# Tropical montane gradients elucidate the contributions of functional traits to competitive and environmental fitness

**DOI:** 10.1101/2023.09.01.555885

**Authors:** Mansi Mungee, Rohan Pandit, Ramana Athreya

**Affiliations:** School of Geography, University of Leeds, Woodhouse lane LS2 9JT, Leeds, UK; Indian Institute of Science Education and Research, Pune Homi Bhabha Road, Pashan, Pune – 411008, Maharashtra, INDIA; B/3 Sterling Homes, Warje, Pune – 411058, Maharashtra

**Author notes:** Corresponding Author Address: Indian Institute of Science Education and Research, Pune Homi Bhabha Road, Pashan, Pune – 411008, Maharashtra, INDIA. MM and RA conceived the ideas and designed methodology; RP collected the data; MM analyzed the data; MM wrote the original draft of the manuscript; MM and RA led the editing and review of the manuscript. All authors gave final approval for publication. We declare that the work presented here is our original research and has not been published elsewhere.

**Keywords:** community assembly, birds, eastern Himalaya, functional traits, phylogenetic diversity

## Abstract

Functional traits can be classified as alpha or beta, based on their relative importance in determining a species’ competitive and environmental fitness, respectively. However, the link between a trait and its contribution to a particular aspect of fitness is not always straight-forward. We investigated phylogenetic and functional diversity for bird communities along a 200-2800 m elevational transect in the eastern Himalayas. We hypothesized that beta traits, associated with environmental tolerances, would exhibit a directional change in mean values, while alpha traits, linked to competitive strategies, would show a decrease in dispersion with elevation. Our findings showed that most functional traits exhibited a decrease in dispersion with elevation. However, surprisingly, the mean values of these traits also exhibited a significant relationship with elevation, suggesting their involvement in both competitive and environmental fitness. Furthermore, we observed that morphological traits, traditionally considered beta traits associated with environmental tolerance, were influenced more strongly by resource availability and habitat structure rather than aspects of temperature or air density. These results challenge the simplistic classification of traits as either alpha or beta. We suggest that future studies should carefully analyze the variation in mean values and dispersion of individual traits before assigning them solely to a particular category of fitness. The results contribute to a broader understanding of the complex interactions between functional traits, fitness, and environmental conditions in Himalayan bird communities.

## 1. Introduction

One of the key goals of tropical ecology is to understand the relative importance of two deterministic assembly processes – environmental filtering and interspecific competition – across taxa and regions (Jarzyna et al. 2021). The large environmental gradients and habitat heterogeneity in montane ecosystems, at relatively small scales, results in a network of local communities which are subsets of the larger ‘regional’ species pool, filtered according to the species’ local environmental requirements . Co-occurring species are further characterized by specialized utilization of resources which minimizes competition and facilitates high local diversity (MacArthur and Levins 1967). The two processes together determine the structure and composition of local communities (Weiher and Keddy 1995).

The presence of these two processes may be inferred by examining the functional and phylogenetic structure of co-occurring species within local communities (Cavender-Bares et al. 2009). Environmental filtering selects for species with similar functional traits that impart adaptive advantages under the local abiotic conditions, resulting in a more clustered or under-dispersed trait composition relative to the regional trait pool (Weiher and Keddy 1995; Swenson and Enquist 2007). Conversely, interspecific competition would tend to segregate functional traits to limit similarity and reduce competition among co-occurring species, resulting in an over-dispersed community trait structure. When these functional traits are phylogenetically conserved – i.e., closely related species have more similar traits – similar patterns of phylogenetic clustering and over-dispersion may also be expected (Coyle et al. 2014).

Lopez et al. (2016) proposed classifying functional traits as alpha and beta according to their perceived importance in determining co-existence and environmental tolerance, respectively (also see Ackerly and Cornwell 2007; Silvertown et al. 2006; Pavoine and Bonsall, 2011). The classification is derived from the concepts of alpha and beta niche (Whittaker 1975), wherein the former determines the division of resources among co-occurring species, while the latter influences a species’ distribution across an environmental gradient. Accordingly, functional traits involved in competitive strategies such as resource acquisition or feeding strategy are classified as alpha traits (e.g. plant height and root depth), and those which determine species’ environmental tolerances, such as the constraints imposed by temperature, oxygen and/or water availability are classified as beta traits (e.g. leaf size and wood density).

The link between a trait and its contribution to a particular aspect of fitness is arguably easier to determine in the case of plants with help of manipulative field and laboratory experiments (Ruprecht et al. 2014; Pérez-Ramos et al. 2019). Not surprisingly, a majority of the studies that use this classification have dealt with plants (Klimeš and Klimešová 2000; Ackerly et al. 2006; Silvertown et al. 2006). Classifying traits in this manner is not always straight-forward with animals, based as it is largely on observations and information from species’ natural histories than on experimental data (Miles and Ricklefs 1984; Pigot et al. 2016). For example, the beak of a bird is commonly considered an alpha trait as it is involved in resource acquisition (Graham et al. 2012). However, the beak, which is uninsulated and well vascularised, is also known to be involved in thermoregulation i.e. environmental tolerance (Tattersall et al. 2017). The relative importance of the beak as an alpha or beta trait will depend on the particular context of the community composition and the environment it inhabits (Friedman et al. 2019). Similarly, body size in birds is commonly considered an important beta trait determining species environmental preferences due to its role in thermoregulation (Gómez et al. 2010). However, disparity in body sizes of co-occurring species is known to promote coexistence (Leyequién et al. 2007); hence it plays a role as an alpha trait as well (Gómez et al. 2010). Many studies of community assembly conflate data from multiple traits to construct a multidimensional picture of species’ niche and/or to improve the statistical signal. This requires a priori knowledge of the association between a trait and the specific fitness category (i.e., alpha or beta) *in the particular study context*.

The change in the community mean value of a trait along the gradient of a particular environmental factor may be considered an indication of an adaptive role of the (functional) trait vis-a-vis that environmental factor (Muscarella and Uriarte 2016). Correspondingly, the dispersion in a trait associated with interspecific competition should increase with competition (Adler et al. 2013). These signatures would be easiest to discern along gradients in interspecific competition and environment since changes in the mean and dispersion are easier to interpret than absolute values (Lopez et al. 2016).

Quite conveniently, elevational transects, especially in the tropics, contain within them both the gradients; it is widely believed that interspecific competition decreases, while environmental filtering increases, towards high elevations (Montaño-Centellas et al. 2021; but see Pérez-Toledo et al. 2022). Harsh abiotic conditions at the higher altitudes, e.g. cold temperatures, limited resources, higher seasonality and low partial-pressure of oxygen, impose stronger environmental filtering, restricting the utility of high elevations to species with traits necessary to cope with their challenges. Conversely, low variability in the abiotic conditions of the lower elevations, coupled with higher productivity and higher species diversity facilitates stronger competitive interactions. Phylogenetic dispersion is arguably a better surrogate of ecological dispersion along such multivariate gradients (in temperature, precipitation, seasonality, oxygen availability, air density, resources, etc.), especially when the extent of a trait’s role in competitive and environmental fitness is not known *a priori* (Webb et al. 2002; Pavoine and Bonsall 2011). This is because phylogenetic dispersion should reflect on the ‘net effect’ of all underlying assembly processes: phylogenetic clustering due to multiple environmental factors and phylogenetic overdispersion due to competition across all conserved traits.

In this study, we first analyzed the variation in phylogenetic dispersion in communities of birds along an east Himalayan elevational transect (200-2800m) to test the expected decline in the relative dominance of interspecific competition, and increase in environmental filtering. We then analyzed the variation in mean and dispersion of several commonly used avian functional traits which have been associated with either one, or both aspects of fitness – environmental and competitive – by previous studies, shown in Table 1. We concluded that a functional trait is: (i) an alpha-trait if the dispersion in the community trait distribution decreases with elevation, and (ii) a beta-trait if its community-mean value exhibits directional variation with elevation

**Table 1.** Functional traits of birds used in the present study to assess variation in community-mean and community-dispersion along an east Himalayan elevational gradient. Classifying functional traits as alpha or beta (according to their perceived importance in determining co-existence and environmental tolerance, respectively) has been suggested to associate patterns in trait dispersion (clustering/overdispersion) with specific community assembly processes (environmental filtering/interspecific competition; see text for more details). However, individual traits may be associated with single, or multiple ecological strategies in birds as shown below.

| Trait | Competitive/environmental fitness | References |
| --- | --- | --- |
| Beak size | 1. Resource-related competition (α-niche)<br>2. Visual and acoustic signalling (α-niche)<br>3. Thermoregulatory function (β-niche) | Gómez et al. 2010; Graham et al. 2012; Luck et al. 2012; Boyce et al. 2019; Pigot et al. 2016 |
| Body-mass | 1. Stress tolerance related to thermoregulation (β-niche)<br>2. Stress tolerance related to recourse availability (β-niche)<br>3. Resource competition (α-niche) | |
| Wing-length | 1. Stress tolerance related to air density (β-niche)<br>2. Stress tolerance related to recourse availability; long-distance dispersal for foraging flights (β-niche)<br>3. Aligned with movement capacity which in turn influences resource use (α-niche) | |
| Tarsus-length | 1. Resource and habitat competition, foraging niche and foraging stratum (α-niche)<br>2. Stress tolerance related to thermoregulation (β-niche) | |
| Primary Substrate | Resource and habitat competition, foraging niche (α-niche) | |
| Foraging Mode |  |  |
| Primary Diet | Resource competition (α-niche) | |
| Habitat | Habitat competition (α-niche) | |

Our study transect is characterized by a steep decline in temperature, resource availability and habitat complexity – all of which may impose environmental stress on ecological communities. The transect also hosts an exceptionally high regional and local bird diversity, and high rates of species turnover (Mungee et al. 2021), providing an ideal system to detect signatures of competitive overdispersion. Further, our fine-scale primary data (50 m elevational resolution) provides a fair picture of ‘local’ communities of co-occurring, and likely interacting species.

## 2. Materials and Methods

### 2.1 Study region, species and trait data

We sampled birds between 200 and 2800 m along a single elevational transect in Eaglenest wildlife sanctuary in the eastern Himalayan global biodiversity hotspot. Birds were recorded visually or aurally along 48 transects of 100 m length, separated from their neighbours by 0.5-2 km of distance and 50 m in elevation, during April-July from 2011 to 2014. A detailed description of the study region and primary data can be obtained from Mungee et al. (2021).

We compiled four species-level morphological traits: beak size (the product of length, width and depth), wing length, tarsus length and body mass. We corrected the (other) morphological traits for allometry with body-mass using linear regression (R^2^ = 0.28-0.89; Supporting Information; Figure S1); i.e. for each morphological trait y,

*residual_y = log_10_(y) – intercept – slope * log_10_(body-mass)*.

We also compiled the species’ behavioral and ecological preferences including primary substrate, foraging mode, primary diet and habitat preferences (Price et al. 2014; Wilman et al. 2014; Schumm et al. 2020; Athreya, 2006; Tobias et al. 2022). All numerical traits were standardized using mean and standard deviation before analyses. Trait-trait correlations were assessed using pairwise Spearman’s correlations.

### 2.2 Phylogenetic data and evolutionary signal

We used Schumm et al.’s (2020) phylogeny of the Himalayan avifauna to estimate evolutionary relationships of species present in our data. The sixteen species absent from this phylogeny were added to their respective clades as polytomies using the global avian phylogeny in Jetz et al. (2012). We used Pagel’s lambda (λ; Pagel 1999) to determine the strength and significance of the phylogenetic signal in each functional trait; i.e. to determine phylogenetic niche conservatism. We chose Pagel’s λ over Blomberg’s K (Blomberg et al. 2003) as it is more robust in the presence of incompletely resolved phylogenies and missing branch-length information (Molina-Venegas and Rodríguez 2017). A value of zero indicates no phylogenetic signal, whereas values ≥ 1 indicate a strong phylogenetic signal (Pagel 1999). We estimated the statistical significance of λ by using likelihood ratio tests which compare the observed trait values to those expected with λ = 0.

### 2.3 Community trait and phylogenetic dispersion along the elevational gradient

We calculated phylogenetic and trait dispersion using Mean Pairwise Distance (MPD) and Mean Nearest Taxon Distance (MNTD; Webb et al. 2002). MPD is the average functional or phylogenetic distance between all co-occurring species in a community. Its value is influenced by the lengths of the branches connecting the deep or ‘basal’ nodes of the functional or phylogenetic tree. On the other hand, MNTD is calculated as the average distance between each species and its closest relative in the local community. Therefore, MNTD is more informative on the dispersion in the ‘terminal’ nodes of the functional/phylogenetic tree. For both metrics, MPD and MNTD, higher values indicate higher trait/phylogenetic dispersion while lower values indicate clustering (Webb et al. 2002).

We calculated MPD and MNTD for (i) each individual functional trait – e.g. MPD_BEAK_ and MNTD_BEAK_ (for beak size), (ii) multivariate functional diversity – MPD_F_ and MNTD_F_, i.e. all the functional traits pooled together, and (iii) phylogenetic diversity – MPD_P_ and MNTD_P_. In the multivariate analysis, we used equal weights for each trait. The functional trait dendrogram (needed to calculate MPD and MNTD) was generated using Gower distances (Gower 1971) since it can accommodate both numerical and categorical traits.

We used null models to determine if the MPD and MNTD metrics differed significantly (over-or under-dispersion) from a random distribution (Webb et al. 2002). Null models that maintain within-community trait-abundance link but randomize habitat occurrences i.e. across community randomization, perform better when estimating under-dispersion in response to environmental filtering. On the other hand, within-community over-dispersion as a result of interspecific interactions are better detected with null models that do not maintain the connection between species abundances and trait values (Gӧtzenberger et al. 2016). tzenberger et al. 2016).

Accordingly, we tested the data using the “*frequency*” null model (*Null-1*) which performs across-habitat randomizations, and the “*independentswap*” null model (*Null-2*) which ignores the link between abundances and traits (Gӧtzenberger et al. 2016). tzenberger et al. 2016). In both cases, we used all species present across all elevations to populate the regional species pool as there are no physical barriers along our study gradient and birds are active dispersers. For more details on the randomization procedures and null models see Supporting Information Table S2.

We calculated Standardized Effect Sizes (SES, Gotelli and McCabe 2002) to quantify the difference between the observed and the null (simulated) values of the metric *X* (MPD or MNTD) using 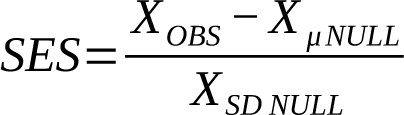, where *μ and SD* are the mean and standard deviation of 999 random simulations of the null model. A negative SES value suggests an under-dispersed community indicative of strong environmental constraints. On the other hand, a positive value suggests over-dispersion which is associated with interspecific competition. We compared the results from null models calculated using species abundance and also presence-absence data using Fisher’s Z-test; the former is expected to increase the sensitivity of detecting under-or over-dispersion (Gӧtzenberger et al. 2016). tzenberger et al. 2016; Tucker et al. 2016). We investigated the relationship between different dispersion metrics and elevation using linear models.

### 2.4 Community trait mean along the elevational gradient

We calculated community-weighted mean (CWM) values for each local community for each trait (Lavorel et al. 2008). For a quantitative trait (relative beak, relative tarsus, relative wing and body mass) CWM is the mean trait value of all species present in the community, weighted by their local abundances. For categorical traits (diet, foraging mode, primary substrate and habitat), CWM is the proportion of each ‘category’ of the trait in the community, i.e. their relative abundance compared to other categories, i.e. functional groups (Lavorel et al. 2008). All calculations were done in the R programming environment (Version 4.3.0; R Foundation for Statistical Computing 2009, http://www.r-project.org/). Pagel’s λ was calculated using the functions “*fitContinuous”* and “*fitDiscrete”* of the R package *Geiger* (Pennell et al. 2014); pairwise species functional distances were computed using the function “*gowdis*” from the package *FD* (Laliberté et al. 2014a; Laliberté et al. 2014b); pairwise species phylogenetic distances were computed with “*cophenetic.phylo*” from package *ape* (Paradis 2010; Paradis et al. 2019); MPD and MNTD metrics, and their SES values were all generated using the package *PICANTE* (Kembel et al. 2010); package *lmtest* was used to compare the slopes of linear models between abundance-based and incidence-based datasets (Hothorn et al. 2015); all CWM values were computed using “*functcomp*” from the package *FD* (Laliberté et al. 2014a; Laliberté et al. 2014b).

## 3. Results

We recorded 15,867 individual birds, spanning 245 species, 150 genera and 50 families. We found strong and significant phylogenetic signal in all functional traits (*λ = 0.78-0.99*) except habitat affinity (*λ = 0.01*), indicating that closely related species possess similar morphology and ecology in our data (Table S1). Standardized effect sizes (SES) for MPD and MNTD were similar across the two null models, Null-1 and Null-2. This was true for each individual trait as well as multivariate functional and phylogenetic metrics. Therefore, in the remaining sections we discuss only the results from Null-1, while results using Null-2 are included in the Supporting Information (Figure S2 - S4).

MPD_F_, but not MNTD_F_, exhibited significant decline with elevation with respect to both observed and SES values (Figure 1). Abundance-based SES-MPD_F_ exhibited a steeper decay with elevation than its incidence-based counterpart (*z = 1.2; p < 0.05*) (Table 2). MPD_P_ and MNTD_P_ were similar to the multivariate functional metrics (Table 2, Figure 1). Mean Pairwise Distances (MPD) exhibited a negative relationship with elevation for all traits, except primary substrate (Table 2, Figure 2 and 3). There were significant differences between the linear regression slopes of abundance- and incidence-based MPD in four traits: relative wing *(Fisher z= -3.72; p < 0.001)*, tarsus *(z = -3.34; p < 0.001),* beak *(z = 4.51; p < 0.001)*, and primary substrate (*z = -3.98; p < 0.001;* Table 2).

**Figure 1.**
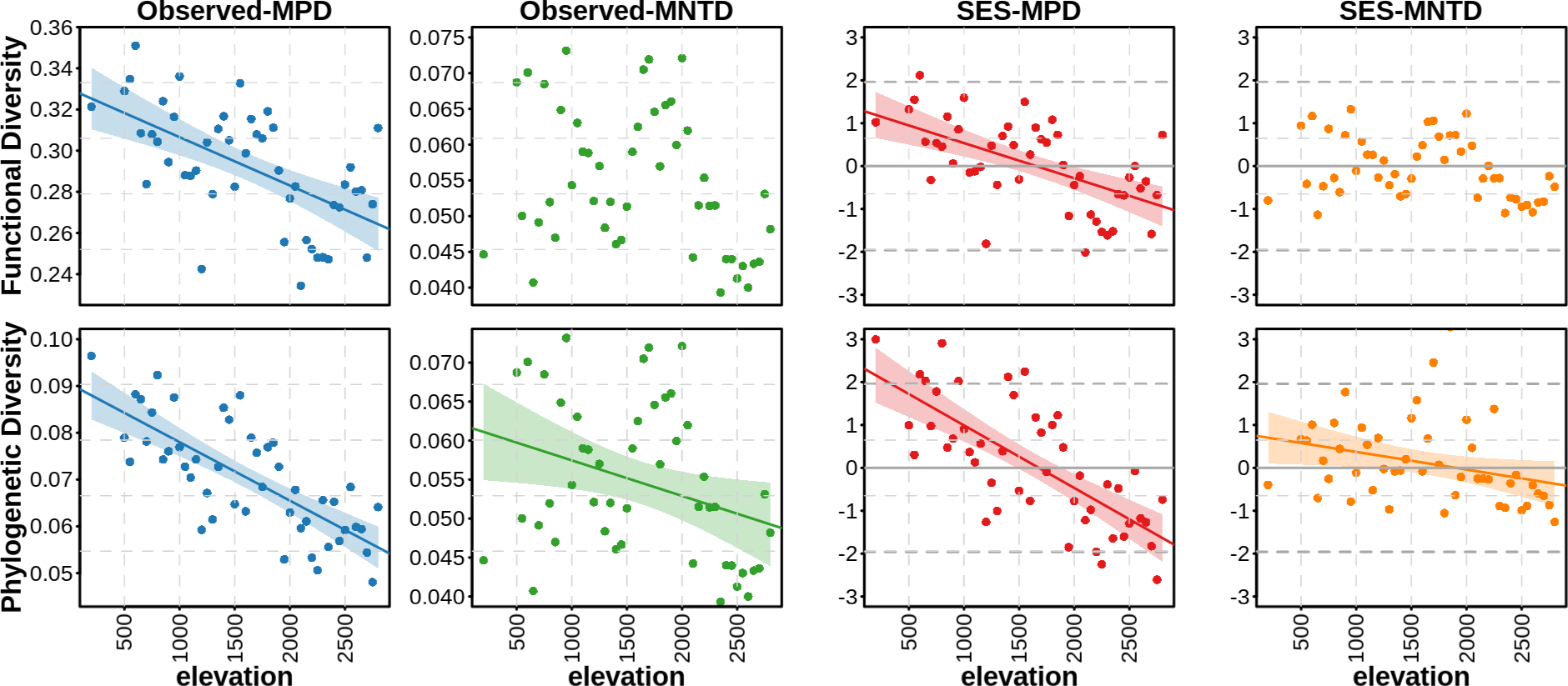
Variation in multivariate functional (all traits pooled) and phylogenetic dispersion for birds along an east Himalayan elevational transect in northeast India. Within each local community, functional and phylogenetic dispersion was measured using two metrics: mean pairwise distance (MPD) and mean nearest taxon distance (MNTD). The observed values for the two metrics (Observed-MPD and Observed-MNTD) were compared against a randomized (null) community using the standardized effect sizes (SES-MPD and SES-MNTD). Within each plot, higher values indicate over-dispersion, while lower values indicate clustering in the community’s trait structure. Significantly clustered assemblages are those with SES values ≤ −1.96, and significantly over-dispersed assemblages were those with SES values ≥ 1.96. However, a significant relationship of dispersion (SES) with elevation is itself an indication of change in clustering/dispersion across the gradient. Fitted linear models are shown only for significant relationships with elevation.

**Table 2.** Variation of community dispersion metrics (MPD and MNTD) of bird functional and phylogenetic diversity along an east Himalayan elevational gradient. Adj. R^2^ (goodness of fit) represents the proportion of variance explained by the model; *p. values* represent the significance of slope (p < 0.01**, p < 0.05*). z-statistic and associated *p. values* are the statistics from Fisher’s Z-test for comparison of abundance versus incidence datasets for the respective trait and metric.

|  | Obs. MPD |  | Obs. MNTD |  | SES-MPD |  | SES-MNTD |  |
| --- | --- | --- | --- | --- | --- | --- | --- | --- |
| | Adj. $R^2$ | *z* | Adj. $R^2$ | *z* | Adj. $R^2$ | *z* | Adj. $R^2$ | *z* |
| Functional (all traits) | 0.34<sup>**</sup> | -0.81 | 0.09<sup>*</sup> | -0.86 | 0.33<sup>**</sup> | 1.20<sup>*</sup> | 0.08<sup>*</sup> | -0.28 |
| Phylogenetic | 0.56<sup>**</sup> | -0.93 | 0.09<sup>*</sup> | -0.86 | 0.52<sup>**</sup> | 2.30<sup>**</sup> | 0.07<sup>*</sup> | -0.66 |
| Relative Wing length | 0.39<sup>**</sup> | -3.72<sup>**</sup> | 0.14<sup>**</sup> | -2.42<sup>**</sup> | 0.34<sup>**</sup> | -2.95<sup>**</sup> | 0.12<sup>*</sup> | -2.16<sup>**</sup> |
| Relative Tarsus length | 0.43<sup>**</sup> | -3.34<sup>**</sup> | 0.01 | -0.37 | 0.39<sup>**</sup> | -1.98<sup>**</sup> | 0.01 | -0.21 |
| Body-mass | 0.28<sup>**</sup> | 1.30 | 0.28<sup>**</sup> | -0.50 | 0.24<sup>**</sup> | 1.08 | 0.26<sup>**</sup> | -1.16 |
| Relative beak size | 0.05 | 4.51<sup>**</sup> | 0.04 | 1.34 | 0.05 | 4.34<sup>**</sup> | 0.04 | 0.66 |
| Primary Substrate | 0.01 | -3.98<sup>**</sup> | 0.12<sup>**</sup> | 0.74 | -0.01 | -4.47<sup>**</sup> | 0.11<sup>*</sup> | -0.25 |
| Foraging Mode | 0.13<sup>**</sup> | 0.16 | -0.02 | 0.80 | 0.12<sup>*</sup> | 2.93<sup>**</sup> | -0.02 | 0.68 |
| Diet | 0.19<sup>**</sup> | 0.93 | -0.20 | 1.59 | 0.20<sup>**</sup> | 2.49<sup>**</sup> | -0.02 | 1.67<sup>**</sup> |
| Habitat | 0.19<sup>**</sup> | 0.94 | -0.02 | 1.59 | 0.20<sup>**</sup> | 2.53<sup>**</sup> | -0.02 | 1.72<sup>**</sup> |

**Figure 2.**
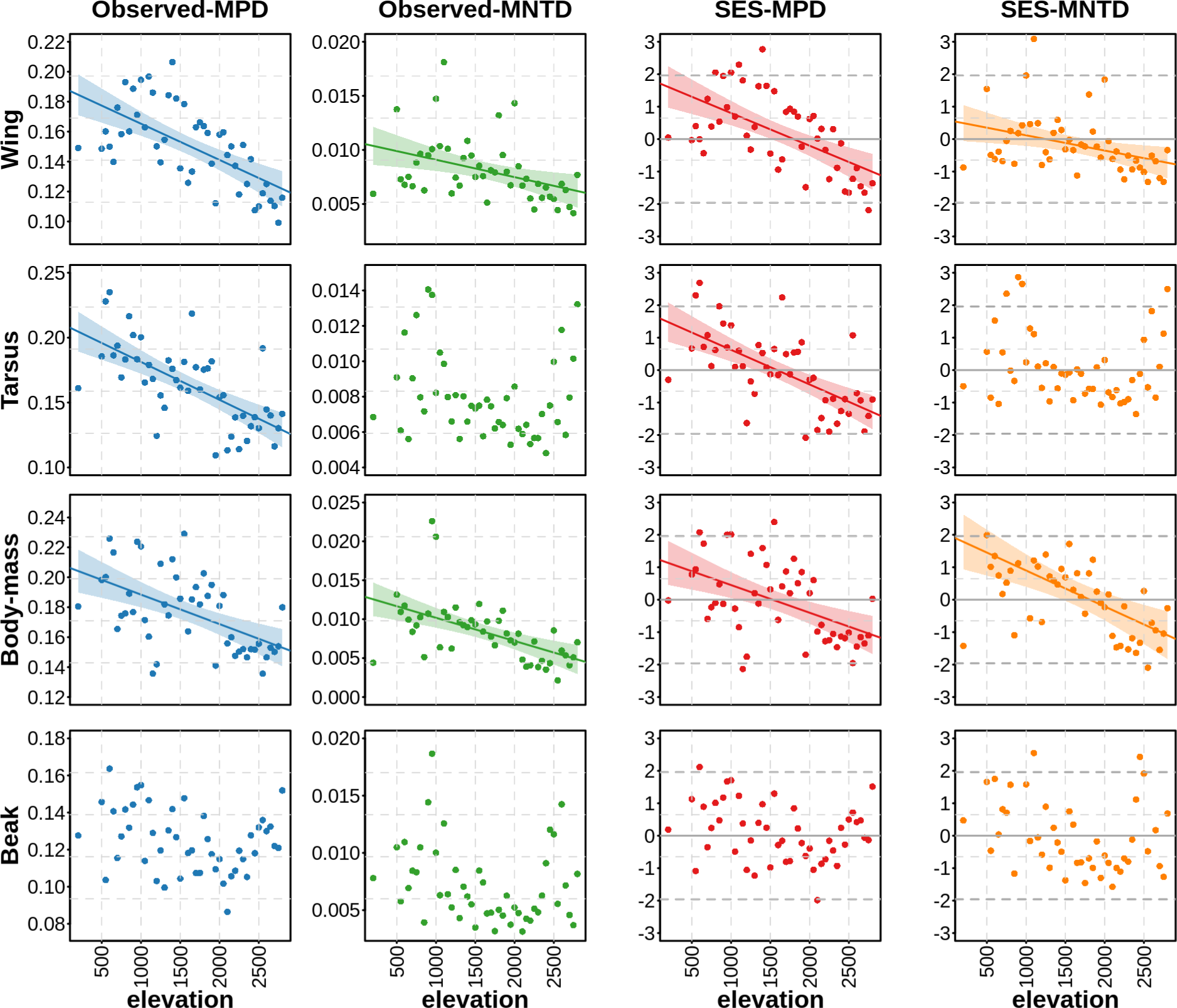
Variation of dispersion in four functional traits (wing and tarsus length, body mass and beak size) for birds along an elevational transect in the Eastern Himalaya of northeast India. Within each local community, dispersion was measured using two metrics: mean pairwise distance (MPD) and mean nearest taxon distance (MNTD). The observed values for the two metrics (Observed-MPD and Observed-MNTD) were compared against a randomized (null) community using the standardized effect sizes (SES-MPD and SES-MNTD). Fitted linear models are shown only for significant relationships with elevation.

The negative relationship with elevation was also seen when MPD was replaced by SES-MPD. We point out that the negative relationships, indicating a change of dispersion with elevation, are significant even though a majority of community-specific SES values lie within ±1.96 SD of the null distribution. The abundance-based metrics exhibited a steeper decay with elevation for all traits except body-mass, habitat, and primary substrate (Table 2). Observed values of MNTD are more variable across the different traits (Table 2, Figure 2 and 3). A significant decline was observed only for relative wing length, beak and body mass. A majority of community SES values lie within ±1.96 SD (Figure 3). Relative tarsus and relative beak size show U-shaped patterns of variation in observed-MNTD and SES-MNTD (Figure 2).

**Figure 3.**
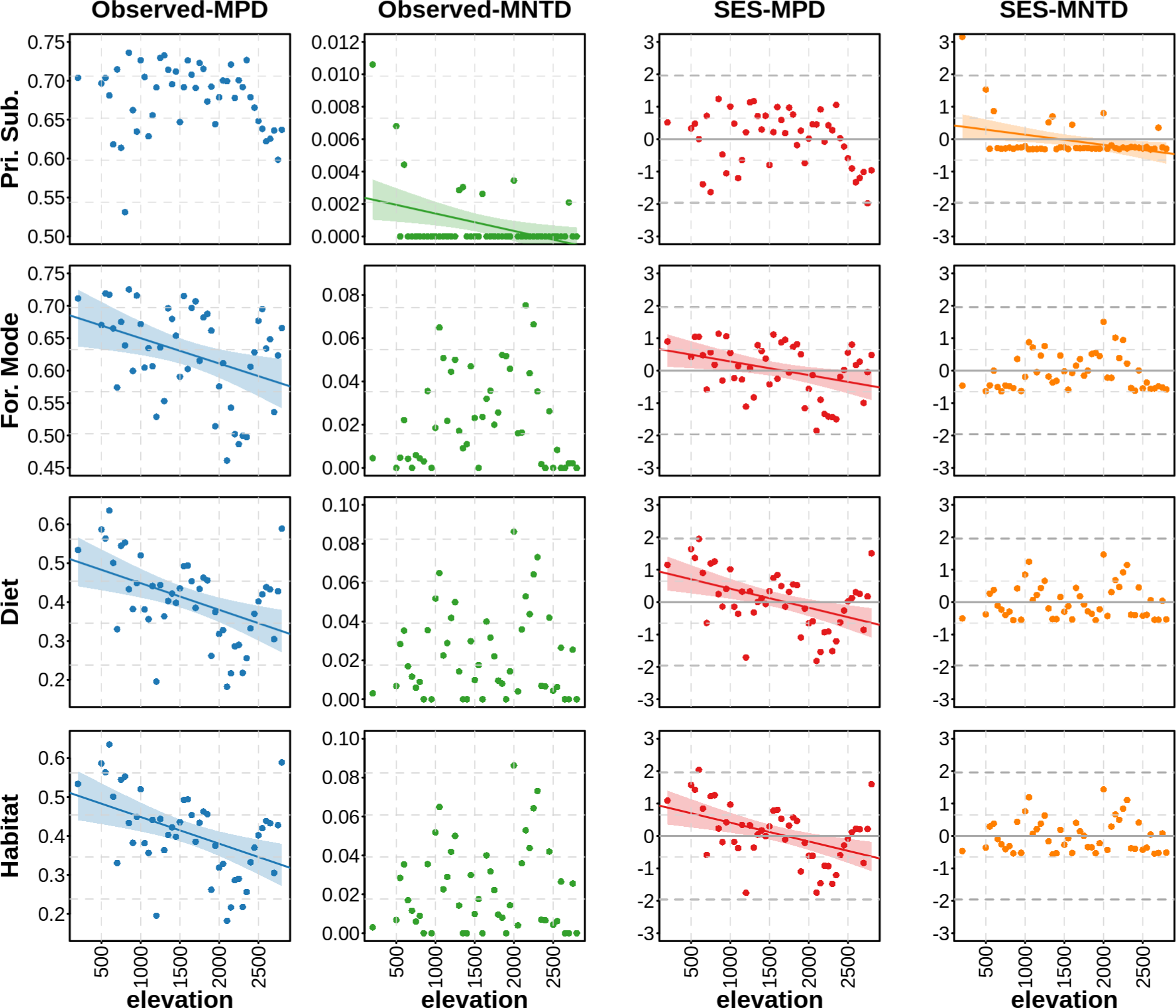
Variation of dispersion in four functional traits (primary substrate, foraging mode, diet and habitat) for birds along an elevational transect in the Eastern Himalaya of northeast India. Within each local community, dispersion was measured using two metrics: mean pairwise distance (MPD) and mean nearest taxon distance (MNTD). The observed values for the two metrics (Observed-MPD and Observed-MNTD) were compared against a randomized (null) community using the standardized effect sizes (SES-MPD and SES-MNTD). Fitted linear models are shown only for significant relationships with elevation.

CWM of relative wing length, relative beak size, and body mass decreased with elevation (R^2^ ranging from 0.29 to 0.71), while that of relative tarsus length increased along the gradient (R^2^ = 0.60). The patterns were less consistent and generally non-linear for the ecological and behavioral (*contra* morphological) traits (Figure 4; Supporting Information Figures S5 – S8). There was a strong reduction in the relative abundance of frugivorous birds, while the insectivores and nectarivores increased towards the highest elevations.

**Figure 4.**
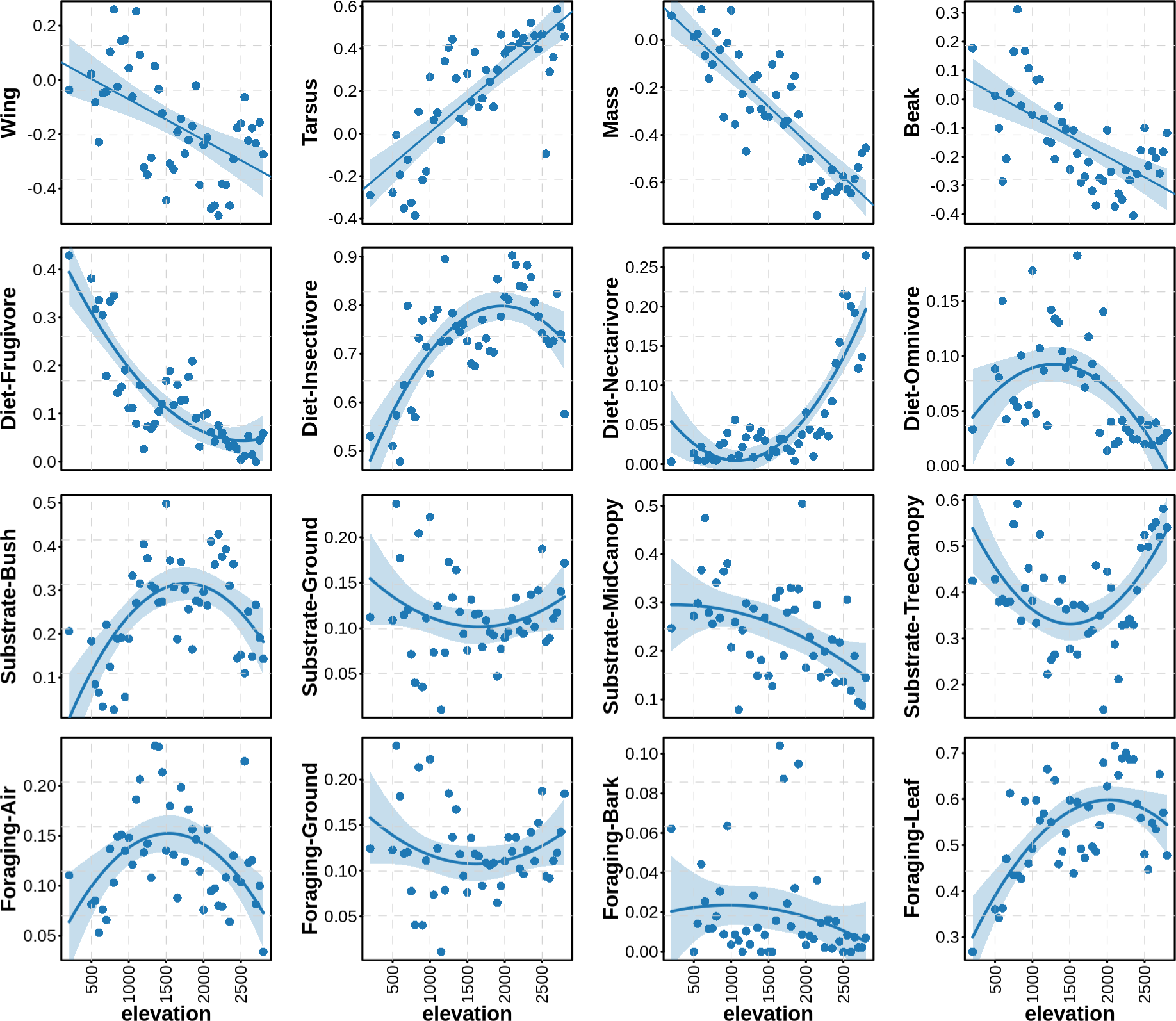
Variation in community weighted mean (CWM) for the different functional traits of birds along an east Himalayan elevational transect in northeast India. CWM values are shown for the four morphological traits in the first row (wing length, tarsus length, body mass and beak size), and the individual categories of different ecological traits in the subsequent rows. For categorical traits (diet, primary substrate and foraging mode, in second, third, and fourth rows, respectively), CWM is the proportion of each functional group in the community, i.e. their relative abundance compared to other functional groups (Lavorel et al. 2008). Wing length, tarsus length and beak sizes were corrected for body-mass allometry prior to the calculation of CWM values (see text for more details). For categorical traits, only four representative categories are shown here. All remaining categories and the remaining ecological trait (Habitat) are included as Supporting Information (Figure S5-S8).

## 4. Discussion

We studied the variation in dispersion and mean of functional traits for bird communities along a 200-2800 m elevational transect in the eastern Himalayas. Along large elevational gradients, environmental stress increases while resource specialization and interspecific competition decreases with elevation. Under these conditions, beta functional traits should exhibit a directional change of their mean value while alpha traits may be identified by a decrease in their dispersion with elevation. We tested this for a set of commonly used functional traits for birds. However, we found that most traits exhibited both trends: their dispersion decreased with elevation, and their mean exhibited a significant relationship with elevation. This suggests that, at least in the context of Himalayan birds, most of the commonly used avian functional traits are implicated in both environmental and competitive fitness.

It is widely believed that interspecific competition decreases while environmental filtering increases towards higher elevations along mountain systems (Graham et al. 2009). This phenomenon, commonly referred to as the Stress-Dominance Hypothesis (SDH), is attributed to the more stressful environment of high elevations which leads to a stabilizing selection of competitively dominant functional strategies, i.e. a convergent or under-dispersed functional trait structure (Weiher and Keddy 1995; Swenson and Enquist 2007; Coyle et al. 2014).

Conversely, the greater specialization needed for sustaining co-existence in the species rich low elevations leads to an over-dispersed trait structure (MacArthur 1969). Conformity to SDH in birds has been observed in a number of studies in tropical mountains (Graham et al. 2009; Dehling et al. 2014; Boyce et al. 2019; Montaño-Centellas et al. 2019; Jarzyna et al. 2021) and for elevational studies across the world (García-Navas et al. 2021; Jarzyna et al. 2021; Montaño-Centellas et al. 2019; He et al. 2018; Ding et al. 2021).

Our results appear to be consistent with the SDH. We found that phylogenetic (and multivariate functional) dispersion decreased with elevation (Figure 1). Our study transect spanning 2600 m in elevation is characterized by a steep decline in temperature, resource availability and habitat complexity – all of which may impose environmental stress on ecological communities. Our spatial sampling (transects 100 m in length, separated from its neighbour by ∼1 km in distance, and 50 m in elevation) defines communities in which the individuals may be assumed to co-occur and compete directly for local resources. In fact, Weiher and Keddy (1995), while formulating Stress Dominance Hypothesis, remarked that over-dispersion is likely to be “restricted to small-scales….where competitive adversity predominates”.

Across all measured dispersion values – phylogenetic, multivariate functional and individual traits – we observed a significant negative relationship between dispersion and elevation using MPD. This relationship was consistently weaker or absent in the case of MNTD (Figures 1-3). The stronger influence of elevation on deeper evolutionary relations (i.e. MPD *contra* MNTD) suggests early radiation and colonization, followed by subsequent emergence of specialization and competitive niche differences. This is consistent with one earlier study from the region which showed that species accumulation in the (eastern) Himalayas is limited by competition for niche space, rather than elevational expansion (Price et al. 2014). Our study reveals that this specialization occurs at much finer scales than previously shown (within 50 m elevational bands). The discussion of the relationship between dispersion and elevation will be confined to MPD in the rest of this paper.

Previous studies have recommended the use of different null models for detecting over- and under-dispersion, especially when simultaneous affects of environmental filtering and interspecific competition are to be expected, as in the present scenario (Gotelli 2000; Gӧtzenberger et al. 2016). tzenberger et al. 2016; Tucker et al. 2016). Typically, the frequency null model has been suggested to capture processes resulting in higher dispersion than the null, while the *independentswap* null model reveals processes related to under-dispersion. Overall, our results were similar across both null models (Figure S2-S4). However, we also note that in all cases the dispersion values lie within ±1.96 S.D. of both the null models. Nevertheless, the reduction in dispersion is both highly significant and consistent across the elevation axis, indicating a gradual but consistent shift in the nature of the dominant assembly process across the elevational transect. We think this strong signal is due to the physical nature of our study transect and data – a smooth, compact but steep environmental gradient sampled at 48 regularly spaced locations. It is quite likely that in the absence of such fine-grain sampling across so many locations the linear correlation would not have been sufficiently strong to compensate for the lack of data outside ±1.96 S.D.

We found that incorporating species abundances in the null models, and in the calculation of the different dispersion metrics improved the strength of the negative slope in most cases (Table 2). Indeed, in a review spanning 2000-2014, Perrone et al. (2017) found that processes related to limiting similarity are better detected with abundance data, although only a few studies have specifically investigated this issue using empirical data (Bernard-Verdier et al. 2012). Our study provide further evidence on the importance of using species abundances when assessing patterns of under- and over-dispersion along environmental gradients in natural communities (also see HilleRisLambers et al. 2012; Münkemüller et al. 2012; Gӧtzenberger et al. 2016). tzenberger et al. 2016). More importantly, our results suggest that the contradictory conclusions among many previous studies may stem, in part, from the use of presence-absence, as opposed to abundance data.

Beak size in birds is most commonly associated with resource-related competition, and therefore alpha-niche (Moermond and Denslow 1985; Dehling et al. 2014). We found a reduction in MPD_BEAK_ with elevation (Figure 2), indicating the expected decrease in competitive interactions for resources. Schoener (1971) observed a similar higher dispersion in beak sizes of tropical insectivorous birds along a latitudinal gradient, and attributed this to the greater diversity of available prey sizes. This suggests beak sizes as important alpha-traits – although not unambiguously, since their community mean value also exhibited a strong decline with increasing elevation (Figure 4). The reduction in beak size is contrary to their expected role in thermoregulation (beta trait; Tattersall et al. 2017), and may be related to the reduction in the size of arthropod prey with elevation (Schumm et al. 2020). We note that 90% of the birds at the highest elevation are insectivores (Figure 4), suggesting resource size, rather than thermoregulatory constraints, driving the beak size variation along the study gradient (Boyce et al. 2019).

The body-mass, and to a lesser extent, wing and tarsus, have previously been considered beta-traits due to their role in stress-tolerance associated with lower temperatures (Gómez et al. 2010). However, the reduction in wing length and body-mass in our study contradicts these expectations (Bergmann’s rule, Allen’s rule) but is in line with previous studies from tropical elevational gradients (Boyce et al. 2019; Schumm et al. 2020). The strong reduction in body size has been attributed to a reduction in the sizes of arthropod preys along the study region (Schumm et al. 2020). The higher elevations in our study region are also characterized by a simpler habitat structure with reduced under-storey which may select for ground foraging species with longer tarsi and smaller relative wings explaining the observed patterns in these traits. Overall, our results indicate a stronger influence of resource and habitat structure, rather than temperature or air density, on all morphological traits. Nevertheless, a strong directional change in their mean value, and a decline in their dispersion suggests their role in both, beta and alpha niches.

Primary substrate, i.e., the principal place a bird obtains its food, forms an important component of species’ behavioral and foraging niche and is a commonly used surrogate for defining interspecific competitive-axis. In our study, we did not find any evidence of reduction in dispersion for primary substrate, although we did observe variation in the dominance structure across the five individual categories. For example, tree-canopy specialists dominated at the lowest and highest elevations, whereas species obtaining their food from bushes peaked at mid-elevations, co-occurring with ground, mid-canopy and tree-canopy species. Other aspects of species behavioral and resource niches tested here, i.e. foraging mode, diet and habitat, exhibited the expected decline in their dispersion with elevation (Figure 3). We also observed an elevational dependence in the mean values of the different categories of these three traits (Figure 4) though the variation is clearly non-linear. Many previous studies have associated morphological traits in birds with in multiple ecological axes. Our results suggest that even ecological and behavioural traits may be implicated as response traits for both competition and environmental filters, via their response to changing environment and habitat which requires a change in the dominant behavioral, and resource acquisition strategy.

## Conclusion

Ecological studies have relied on natural history observations to associate some of the commonly measured functional traits with either competitive (alpha) or environmental (beta) fitness. Our study indicates that this dichotomy is an oversimplification. While the global associations between avian morphology and ecological function are widely acknowledged— for instance, the beak serving as the primary tool for capturing and processing food, and wings, tails, and legs being linked to locomotion—the extent to which these functions contribute to competition or environmental tolerance is contingent upon the specific circumstances of the species and its environment. We argue that unambiguous classification of functional traits, exclusively associated with either alpha or beta niches, is not straightforward. In future studies employing such classifications, it is crucial to rigorously examine patterns of variation in the mean and dispersion of individual traits before assigning a trait solely to a specific aspect of fitness.

## Acknowledgements

We are grateful for the enormous support from Singchung Bugun community, especially Mr. Indi Glow, Nima Tsering and Puspalal Sharma, and Forest Department of Arunachal Pradesh

## Declarations

**Funding:** The field work of this project was carried out with financial support from the Department of Science and Technology, Government of India (Grant No. SR/SO/AS66/2011), the Nadathur Trust, Bengaluru and IISER, Pune.

## Ethics approval

Ethics approval is not applicable in this study

## Consent to participate

Not applicable

## Consent for publication

Not applicable

## Availability of data and material

The datasets used during the current study will be made publicly available on Dryad upon acceptance.

## Code availability

Publicly available software and packages were used in this manuscript and have been adequately mentioned and cited throughout.

## Authors’ contributions

MM and RA conceived the ideas and designed methodology; RP collected the data; MM analyzed the data; MM wrote the original draft of the manuscript; MM and RA led the editing and review of the manuscript. All authors gave final approval for publication.

