## Supporting Information for "Tropical montane gradients elucidate the contributions of functional traits to competitive and environmental fitness"

**Table S1. Phylogenetic signals (Pagel's lambda) in bird functional traits used in this study.** For each trait, Pagel's lambda (**λ)** was calculated to show influence of phylogeny on trait variation. Lambda values vary from 0 (no influence of phylogeny) to 1 (strong phylogenetic influence). *p-values* correspond to a comparison of the log-likelihoods of a model with the maximum likelihood estimate of lambda for a given trait to the log-likelihood of a model where lambda was set to zero. ($p < 0.01$**, $p < 0.05$* )

| **Trait** | **λ** |
|---|---|
| Relative wing size | 0.79** |
| Relative tarsus length | 0.92** |
| Body-mass | 0.90** |
| Relative beak size | 0.78** |
| Primary substrate | 0.99** |
| Foraging height | 0.98** |
| Primary Diet | 0.93** |
| Habitat | 0.01** |

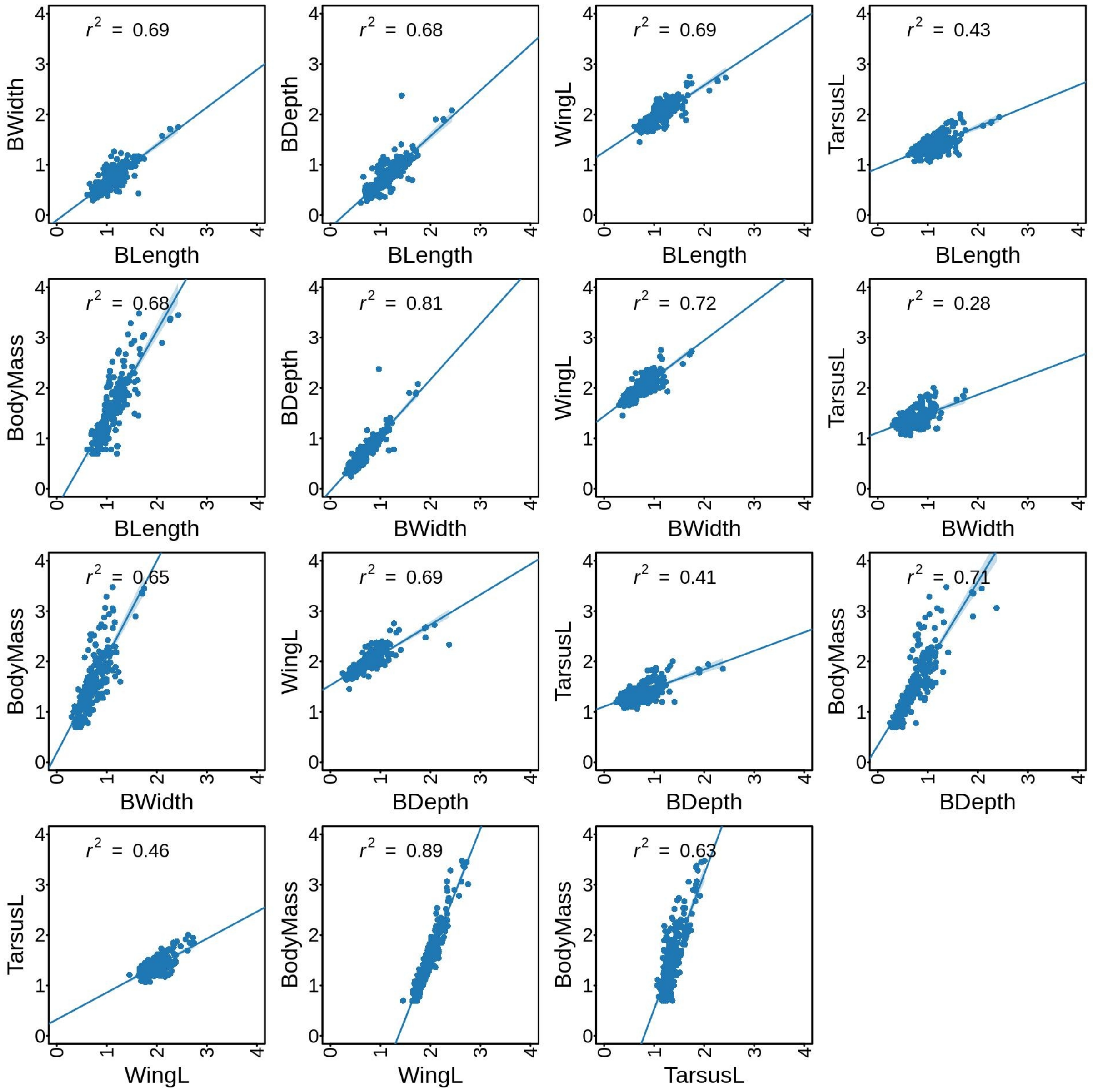


**Figure S1.** Pairwise correlations among all morphological traits for bird species present in our data. The values in the topleft corner of each plot shows the Spearman's correlation coefficients. All traits were log10 transformed before correlation analyses.

**Table S2. Overview of randomizations assessed in this study and their constraints on the community abundance, species richness and occurrence frequency.**

| | **Total Species Abundance Fixed** | **Species Occurence Frequency Fixed** | **Local community Richness Fixed** | **Local commmunity's total abundance Fixed** | **R function and package used** |
|---|---|---|---|---|---|
| **Null-1** *sensu* "C4" null model in Götzenberger et al. (2016) | √ | √ | **X** | **X** | ses.mpd/ ses.mntd from picante with "frequency" randomization |
| **Null-2** | **X** | √ | √ | **X** | ses.mpd/ ses.mntd from picante with "independentswap" randomization |

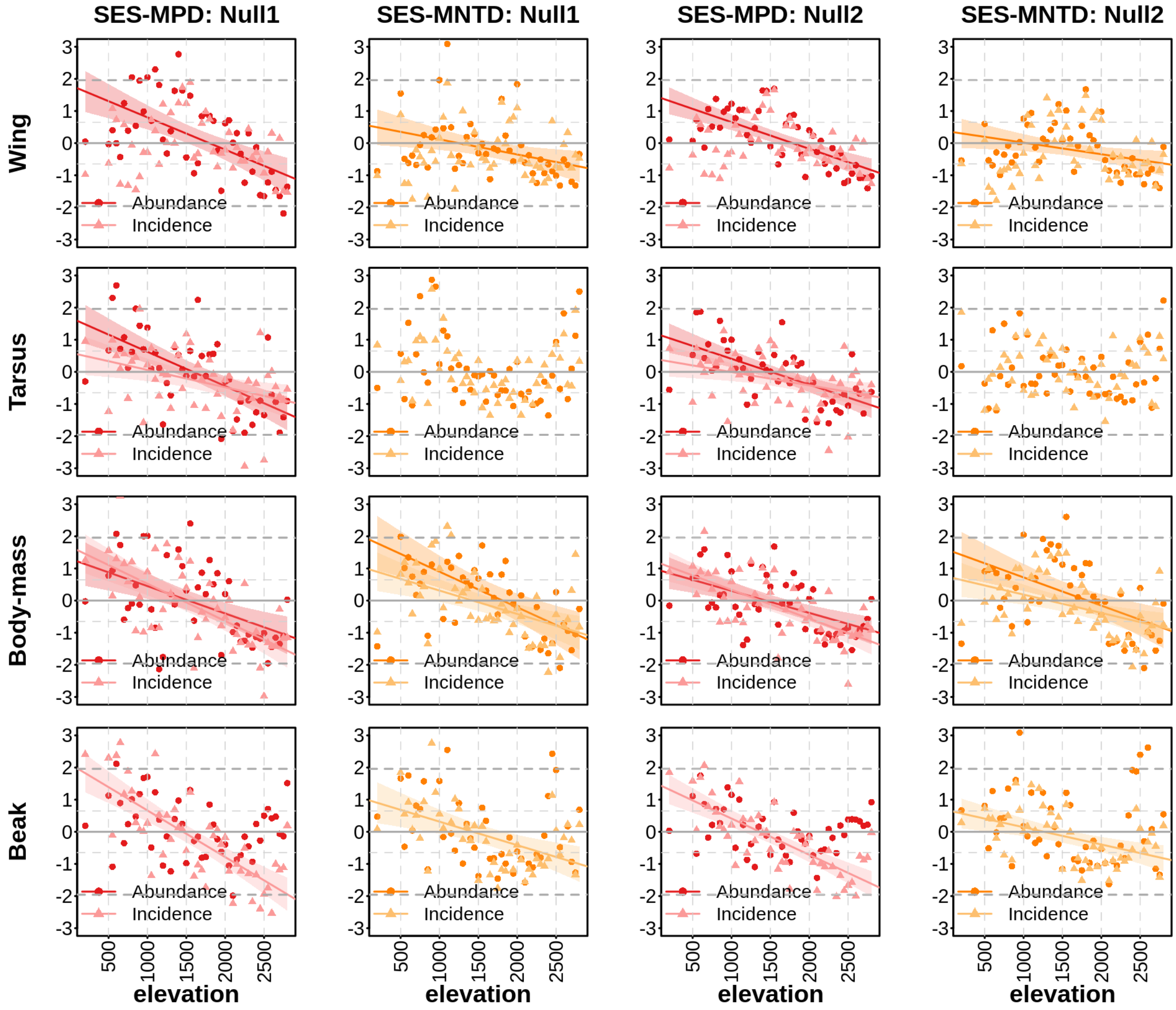


**Figure S2. Variation of dispersion in four functional traits (wing and tarsus length, body mass and beak size) for birds along an elevational transect in the Eastern Himalaya of northeast India.** Within each local community, dispersion was measured using two metrics: mean pairwise distance (MPD) and mean nearest taxon distance (MNTD). The observed values (shown in Main Figure 2) for the two metrics were compared against a randomized (null) community using the standardized effect sizes (SES-MPD and SES-MNTD). We used two different null models: "*frequency*" null model (Null1) which performs across-habitat randomizations, and the "*independentswap*" null model which releases the link between abundances and traits (Null2). SES values represent the deviation of observed metrics from the mean of the distribution of null-values, standardized by the standard deviation of the null-values (see main text for more details). Within each plot, higher (positive SES) values indicate over-dispersion, while lower (negative SES) values indicate clustering in the community's trait structure. Fitted models are shown only for significant relationships. Within each plot, darker colored dots and lines represent dispersion metrics calculated using species relative abundances, whereas lighter colored triangles and lines are based on incidence based data, i.e. using only species occurrences.

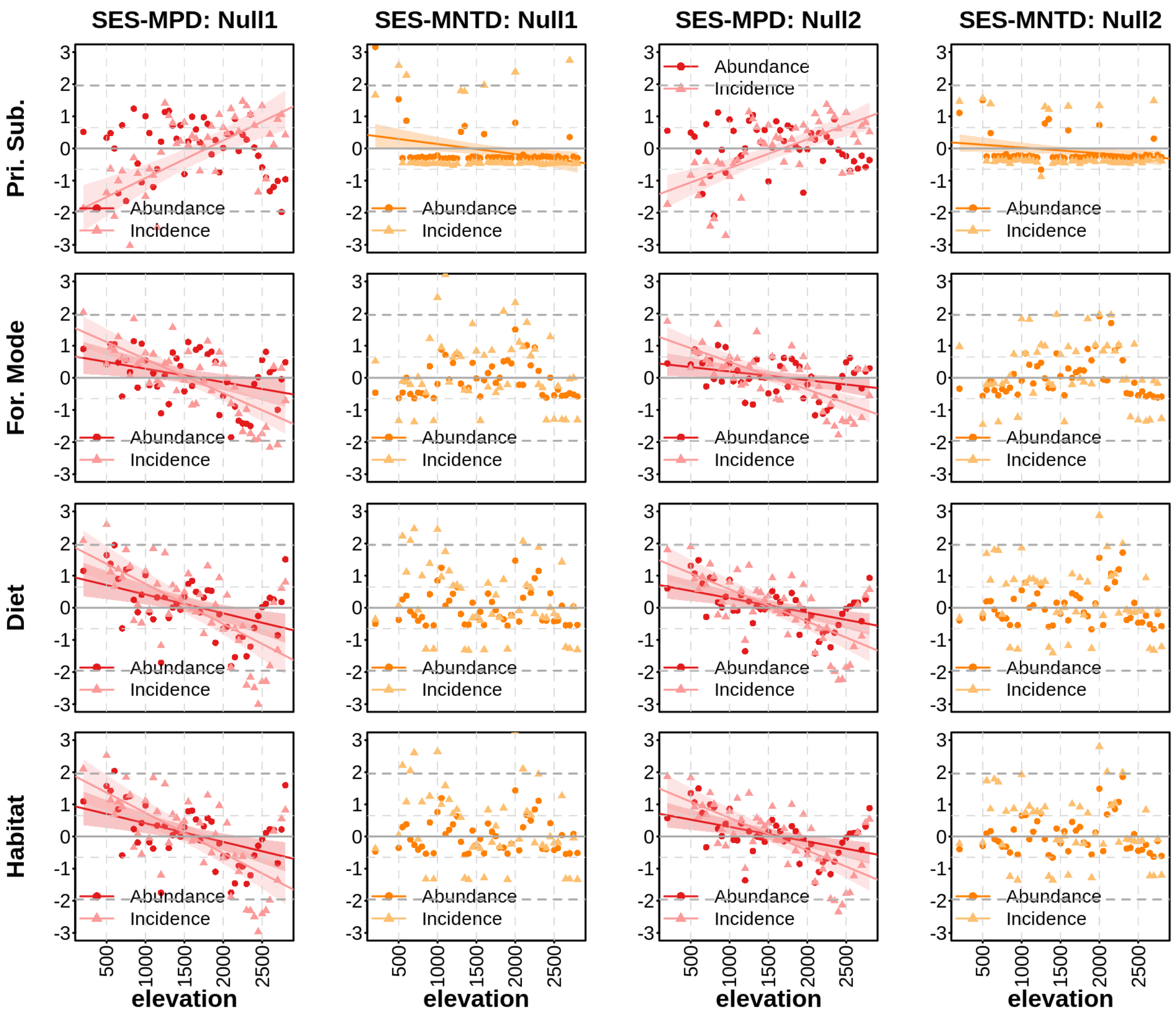


**Figure S3. Variation of dispersion in four functional traits (primary substrate, foraging mode, diet and habitat) for birds along an elevational transect in the Eastern Himalaya of northeast India.** Within each local community, dispersion was measured using two metrics: mean pairwise distance (MPD) and mean nearest taxon distance (MNTD). The observed values (shown in Main Figure 3) for the two metrics were compared against a randomized (null) community using the standardized effect sizes (SES-MPD and SES-MNTD). We used two different null models: "*frequency*" null model (Null1) which performs across-habitat randomizations, and the "*independentswap*" null model which releases the link between abundances and traits (Null2). SES values represent the deviation of observed metrics from the mean of the distribution of null-values, standardized by the standard deviation of the null-values (see main text for more details). Within each plot, higher (positive SES) values indicate over-dispersion, while lower (negative SES) values indicate clustering in the community's trait structure. Fitted models are shown only for significant relationships. Within each plot, darker colored dots and lines represent dispersion metrics calculated using species relative abundances, whereas lighter colored triangles and lines are based on incidence based data, i.e. using only species occurrences.

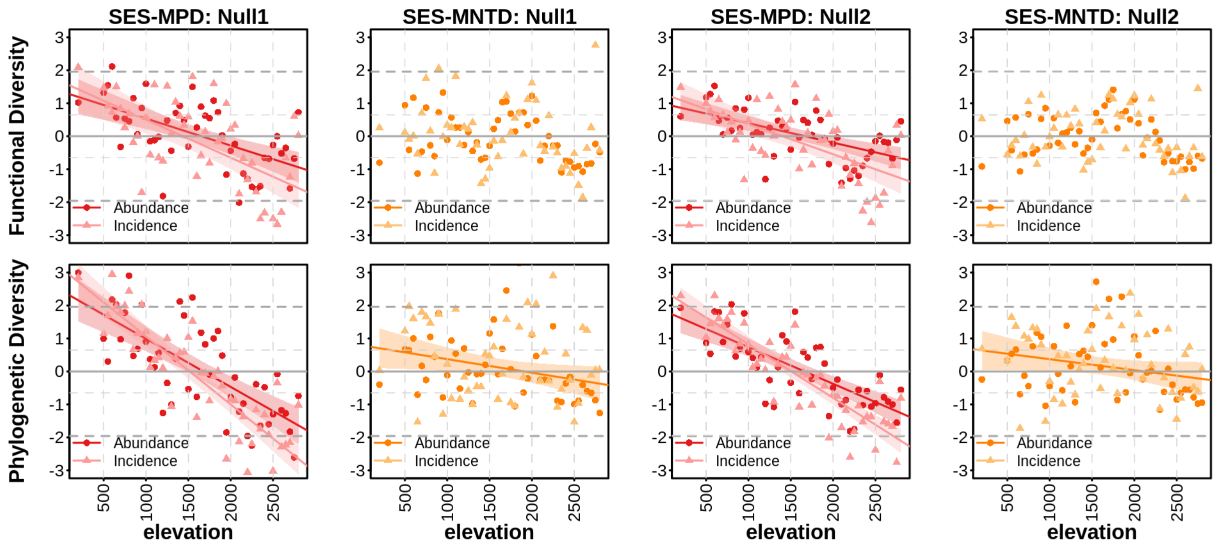


**Figure S4. Variation in multivariate functional (all traits pooled) and phylogenetic dispersion for birds along an east Himalayan elevational transect in northeast India.** Within each local community, functional and phylogenetic dispersion was measured using two metrics: mean pairwise distance (MPD) and mean nearest taxon distance (MNTD). The observed values (shown in Main Figure 4) for the two metrics were compared against a randomized (null) community using the standardized effect sizes (SES-MPD and SES-MNTD). We used two different null models: "*frequency*" null model (Null1) which performs across-habitat randomizations, and the "*independentswap*" null model which releases the link between abundances and traits (Null2). SES values represent the deviation of observed metrics from the mean of the distribution of null-values, standardized by the standard deviation of the null-values (see main text for more details). Within each plot, higher (positive SES) values indicate over-dispersion, while lower (negative SES) values indicate clustering in the community's trait structure. Within each plot, darker colored dots and lines represent dispersion metrics calculated using species relative abundances, whereas lighter colored triangles and lines are based on incidence based data, i.e. using only species occurrences.

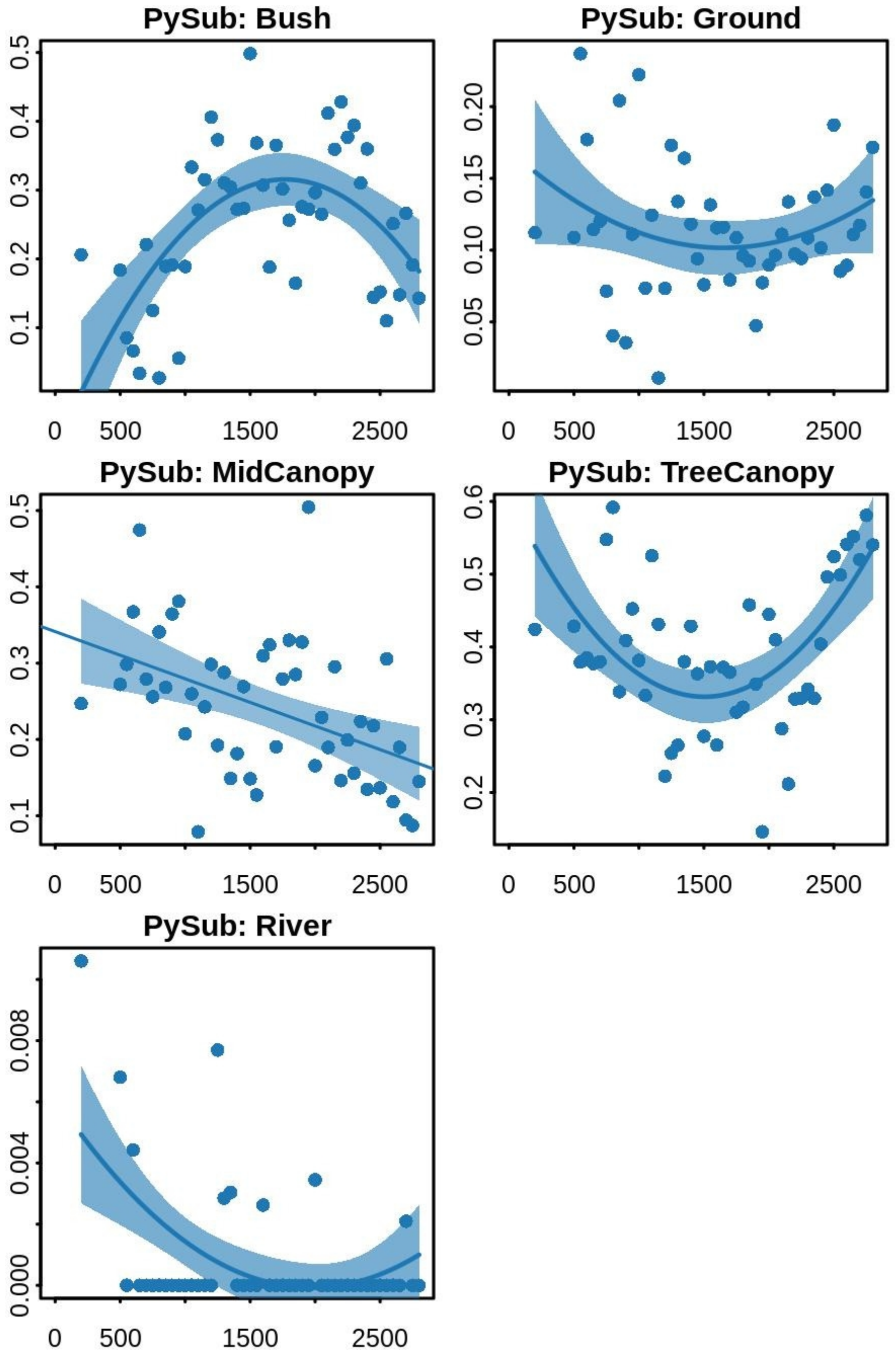


**Figure S5.** Variation in community weighted mean (CWM) for different categories of the ecological trait "Primary Substrate" for birds along the studied elevational gradient. The plot represents CWMs were calculated using the function "*dbFD*" from R package FD 1.0-12 (Laliberté & Legendre 2010; Laliberté et al. 2004; R Core Programming Version 4.1.1). The CWMs represent the abundance of each individual category along the elevational gradient (when CWM.type is "all" in the function *functcomp*). Categories "moss" and "river" not shown due to very few number of species in these categories

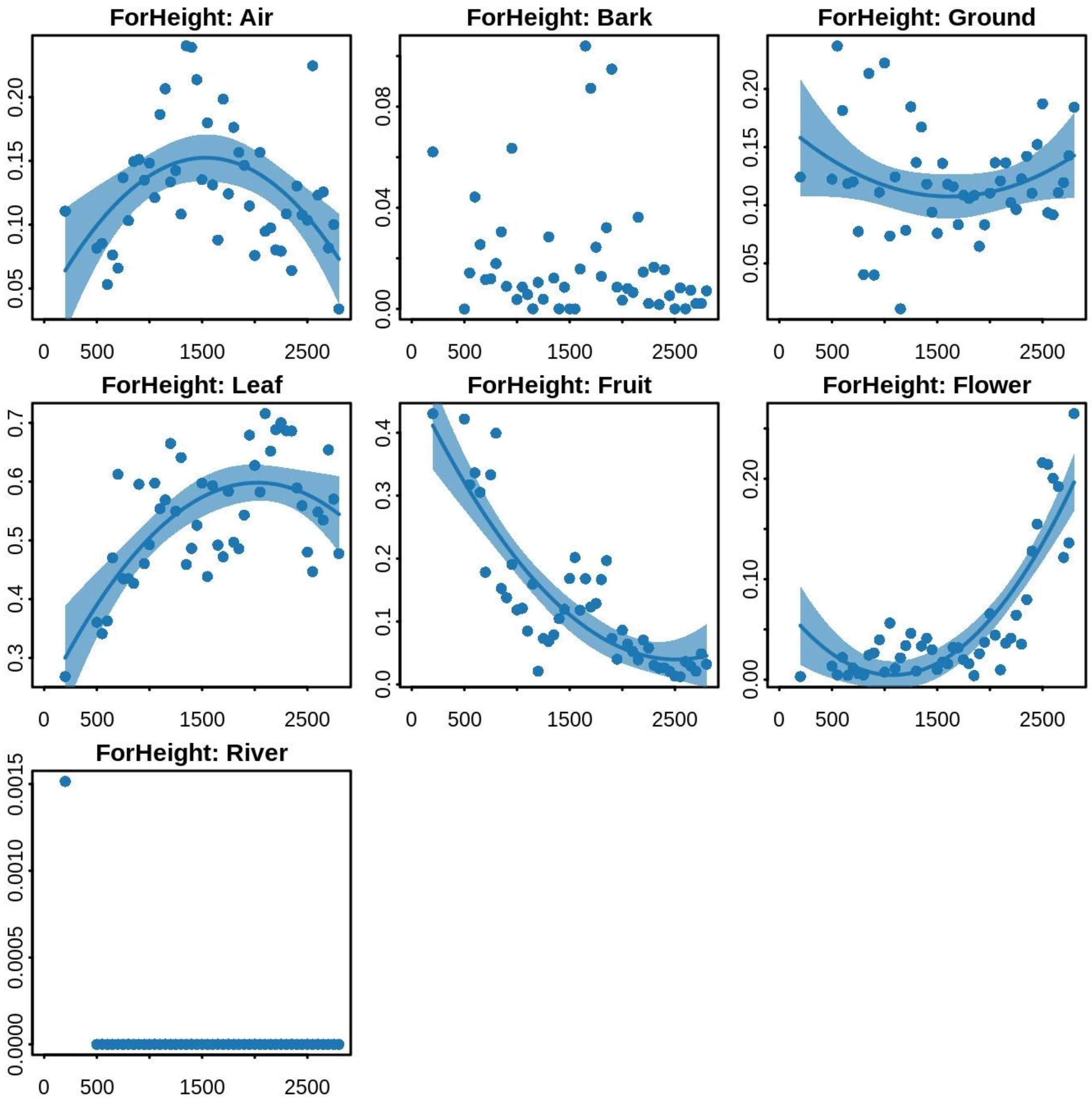


**Figure S6.** Variation in community weighted mean (CWM) for different categories of the ecological trait “Foraging Height” for birds along the studied elevational gradient. The plot represents  CWMs were calculated using the function “*dbFD*” from R package FD 1.0-12 (Laliberté & Legendre 2010; Laliberté et al. 2004; R Core Programming Version 4.1.1). The CWMs represent the abundance of each individual category along the elevational gradient (when CWM.type is "all" in the function *functcomp*). Categories “moss” and “river” not shown due to very few number of species in these categories

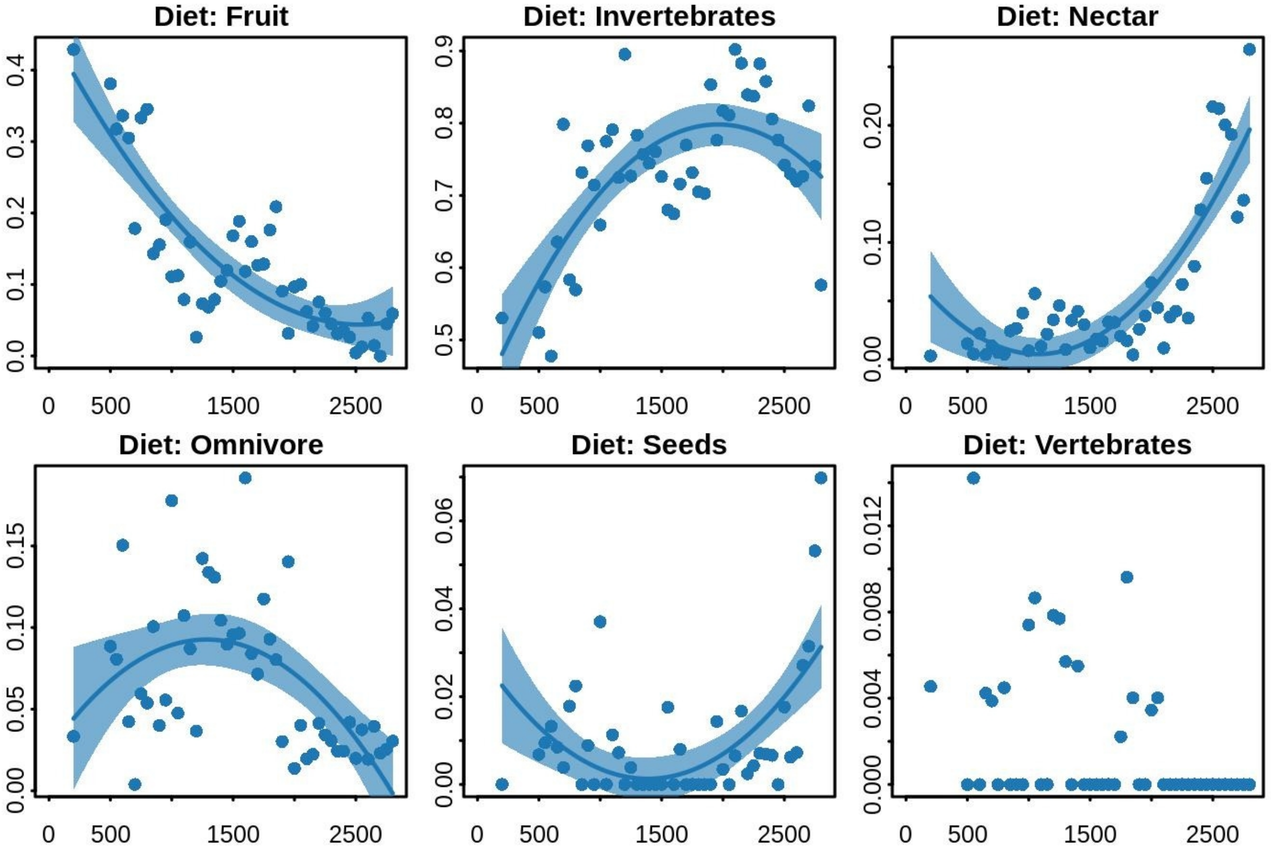


**Figure S7.** Variation in community weighted mean (CWM) for different categories of the ecological trait “Diet” for birds along the studied elevational gradient. The plot represents CWMs were calculated using the function “*dbFD*” from R package FD 1.0-12 (Laliberté & Legendre 2010; Laliberté et al. 2004; R Core Programming Version 4.1.1). The CWMs represent the abundance of each individual category along the elevational gradient (when CWM.type is "all" in the function *functcomp*). Categories “moss” and “river” not shown due to very few number of species in these categories

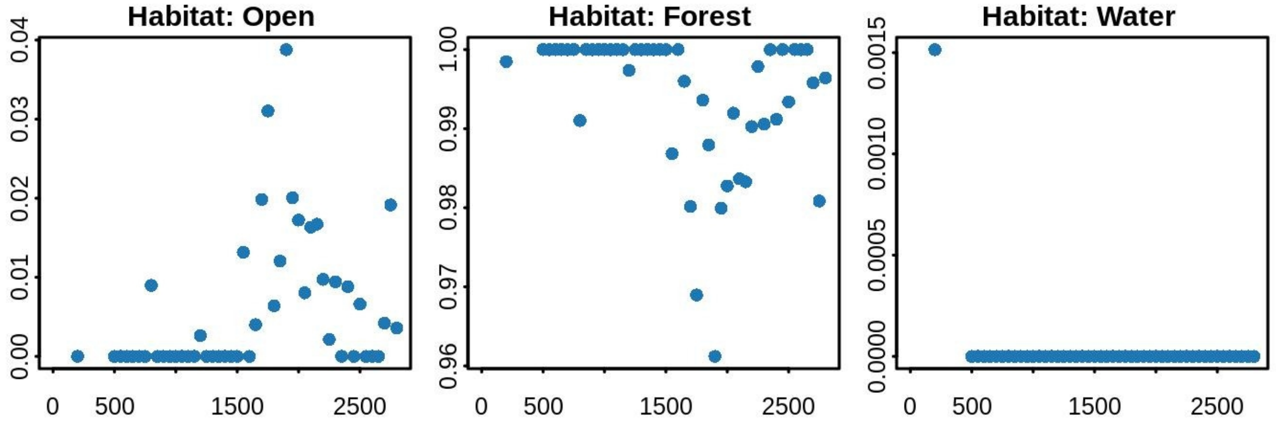


**Figure S8.** Variation in community weighted mean (CWM) for different categories of the ecological trait "Habitat" for birds along the studied elevational gradient. The plot represents  CWMs were calculated using the function "*dbFD*" from R package FD 1.0-12 (Laliberté & Legendre 2010; Laliberté et al. 2004; R Core Programming Version 4.1.1). The CWMs represent the abundance of each individual category along the elevational gradient (when CWM.type is "all" in the function *functcomp*).
